# TaoChongBao: A Large-Scale *Caenorhabditis elegans* Mutagenesis and Missense Variant Database Bridging Worm and Human Genomes

**DOI:** 10.64898/2025.12.03.692224

**Authors:** Ming Li, Shimin Wang, Yongping Chai, Zhengyang Guo, Zi Wang, Zhe Chen, Kexin Lei, Jingyi Ke, Xingshen Huang, Kaiming Xu, Zijie Shen, Wei Li, Guangshuo Ou

**Author notes:** Co-first authors (Ming Li, Shimin Wang).

## Abstract

We generated and sequenced 12,069 viable *Caenorhabditis elegans* strains produced by ethyl methanesulfonate mutagenesis, identifying 20,315,536 variants, including 541,102 unique missense mutations across 20,914 genes. To organize and visualize this resource, we developed TaoChongBao, an open-access database and strain repository that integrates *C. elegans* mutation data with AlphaMissense-predicted pathogenicity scores and ClinVar clinical annotations. TaoChongBao enables users to explore worm missense variants, identify conserved residues corresponding to human pathogenic sites, and access viable strains for experimental validation. Compared with the previous Million Mutation Project in *C. elegans*, TaoChongBao expands mutation coverage over twenty-fold and emphasizes amino acid-altering variants. This resource provides a scalable platform for functional residuomics, variant interpretation, and comparative analyses between *C. elegans* and human genomes.

## Introduction

Single-nucleotide substitutions that change one amino acid—known as missense mutations—can profoundly influence protein structure, stability, and interaction networks. Such substitutions constitute a major class of genetic variation in both health and disease. Understanding the functional consequences of missense mutations is therefore a fundamental goal of molecular biology and precision medicine (1–5). Recent computational advances, such as AlphaMissense, have predicted the pathogenic potential of all 71 million possible single amino acid substitutions in the human proteome. However, translating these predictions into biological understanding requires scalable experimental models that can evaluate the effects of these substitutions in vivo (2). Current approaches, such as CRISPR-based genome editing, allow precise generation of individual missense variants but remain time-consuming and cost-intensive, typically producing one mutation at a time (6,7). A complementary strategy is needed to generate, catalog, and experimentally access large numbers of missense mutations efficiently.

Chemical mutagenesis using ethyl methanesulfonate (EMS) provides such a high-throughput route. Since Sydney Brenner’s pioneering work in the 1960s, EMS mutagenesis has been a cornerstone of *Caenorhabditis elegans* genetics (8–10). Each mutagenized worm typically carries hundreds of single-nucleotide changes, including dozens of missense variants (11–13), and the simplicity of nematode culture allows the generation of vast mutant populations at minimal cost (10,14). With recent advances in high-throughput whole-genome sequencing, it has become feasible to systematically identify and analyze these variants across thousands of strains (13,15–17).

Here, we present a large-scale *C. elegans* EMS mutagenesis and sequencing initiative comprising more than 12,069 strains and yielding 20,315,536 variants—about twenty times the scale of the Million Mutation Project (13). This includes 541,102 unique missense mutations across nearly all coding genes. We developed TaoChongBao, a public database integrating this mutation resource with AlphaMissense predictions (2) and ClinVar annotations (4), enabling users to query conserved worm–human missense sites, visualize pathogenicity information, and access strains carrying relevant variants. Beyond its scale, several features distinguish this resource. Despite heavy mutational loads, all strains remain viable, revealing broad tolerance to single–amino acid substitutions in essential genes and providing insight into mutational robustness. Our screen also incorporated selection with ivermectin, enriching for mutants with ciliary or ion channel defects relevant to neurobiology and drug response (18–21). By mapping conserved residues between worm and human proteins, this dataset offers an experimental platform to interpret human variants of uncertain significance. Together, these advances establish *C. elegans* as a scalable and economical model for functional residuomics and variant interpretation.

## Results

### Large-scale EMS mutagenesis and whole-genome sequencing of *C. elegans* strains

Following an established EMS mutagenesis protocol (Fig. 1A, details in Materials and Methods), we generated 12,069 viable *C. elegans* strains through successive rounds of mutagenesis, selection, and isolation. Notably, 78.4% of these strains exhibit a diverse array of phenotypes, including dumpy morphology, uncoordinated movement, multivulva formation, blistered cuticle and most commonly, resistance to the anti-nematode drug ivermectin (18–21) (Fig. 1B).

**Fig. 1.**
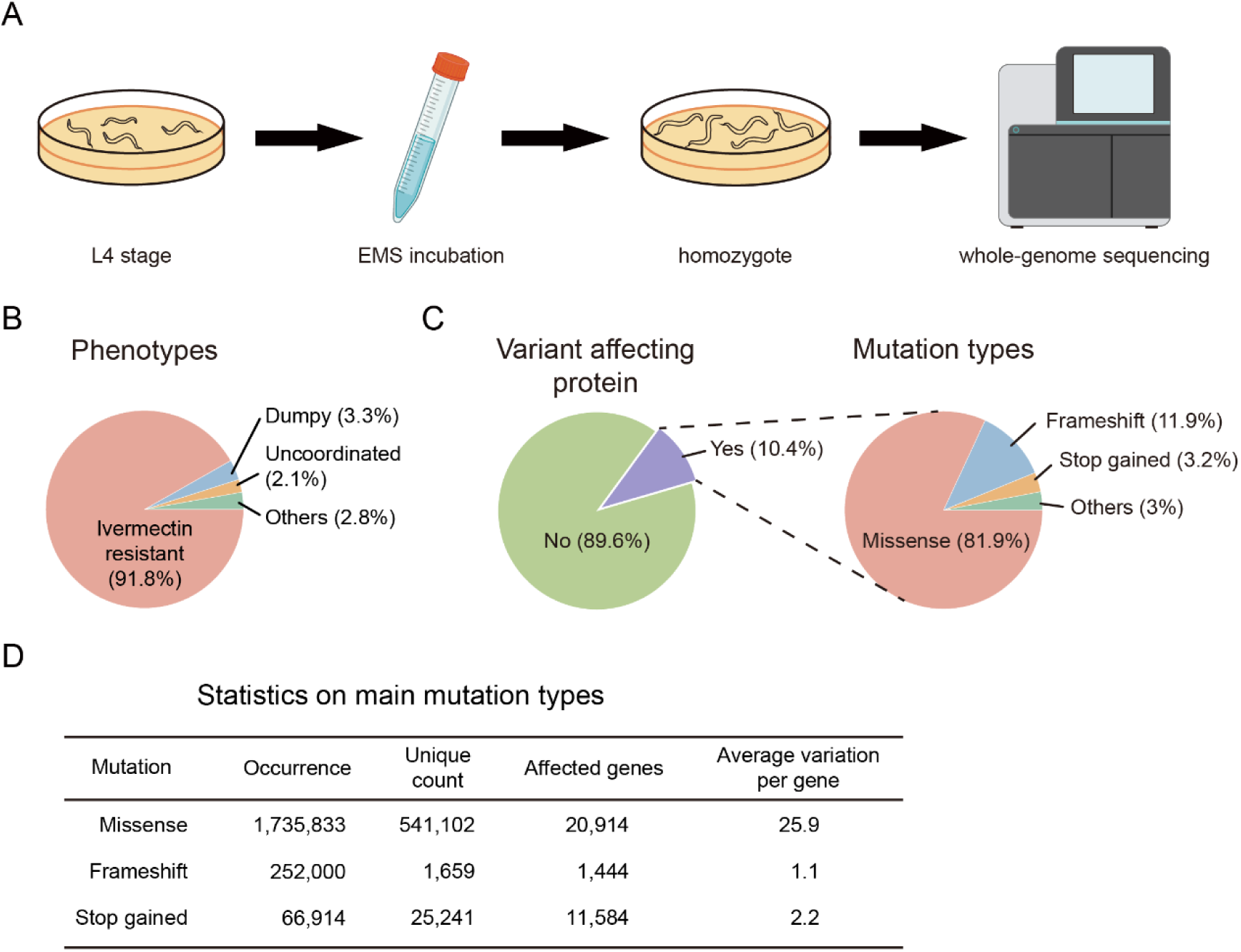
Generation of *C. elegans* mutant collection. (A) Flowchart depicts the generation of *C. elegans* mutants. Late L4 animals were mutagenized with EMS. F2 progenies were singled and propagated, and F3 homozygous mutants were selected for whole-genome sequencing. (B) The phenotypes observed in the *C. elegans* mutant collection. (C) Mutation types identified in the *C. elegans* mutant collection. (D) Statistics for the three major classes of mutations.

Whole-genome sequencing (WGS) was performed for all these strains, achieving an average coverage depth exceeding 15× per genome. Across all strains, we identified 20,315,536 variants distributed throughout the *C. elegans* genome. Consistent with previously reported EMS mutagenesis rates (11–13), each strain harbored an average of ∼1,700 variants, including approximately 144 missense mutations that affect protein-coding sequences. Among all detected variants, 89.6% were located in non-coding regions—including introns, untranslated regions, and intergenic sequences, while 10.4% resided within protein-coding exons. Within the coding subset, 81.9% were missense mutations, followed by frameshift (11.9%), nonsense (3.2%), and other categories such as splice site and stop-loss mutations (3%) (Fig. 1C). In total, 541,102 unique missense mutations were mapped across 20,914 *C. elegans* genes, corresponding to an average of 26 distinct amino acid substitutions per gene (Fig. 1D). Notably, all strains remained viable, implying that most single amino acid substitutions, even in essential genes, do not compromise viability, thus offering a broad collection of functionally tolerant protein variants.

### Construction of the TaoChongBao database

To effectively organize and leverage this extensive genetic dataset, particularly as a resource for genetic disease research, we developed the TaoChongBao database (Fig. 2). First, we retrieved *C. elegans*-human ortholog pairs from the OrthoList 2 project (22), obtaining 28,298 pairs, and supplemented them with an additional 4,572 pairs manually clustered by MMseqs2 (23). We then collected protein sequences for the corresponding *C. elegans* genes from WormBase (24), and human genes from UniProt (25). Recognizing the probability that multiple protein isoforms can be encoded by single gene in both species, we retained all *C. elegans* isoforms presented in our sequencing data, whereas for human genes we retained only the isoform designated as “canonical” by UniProt. This selection yielded 13,790 *C. elegans* isoforms and 11,137 human isoforms. Subsequently, we aligned each isoform pair using Clustal W package with the BLOSUM62 scoring matrix (26), revealing nearly 3,690,301 conserved amino acid residues between *C. elegans* and human proteins (Fig. 3A).

**Fig. 2.**
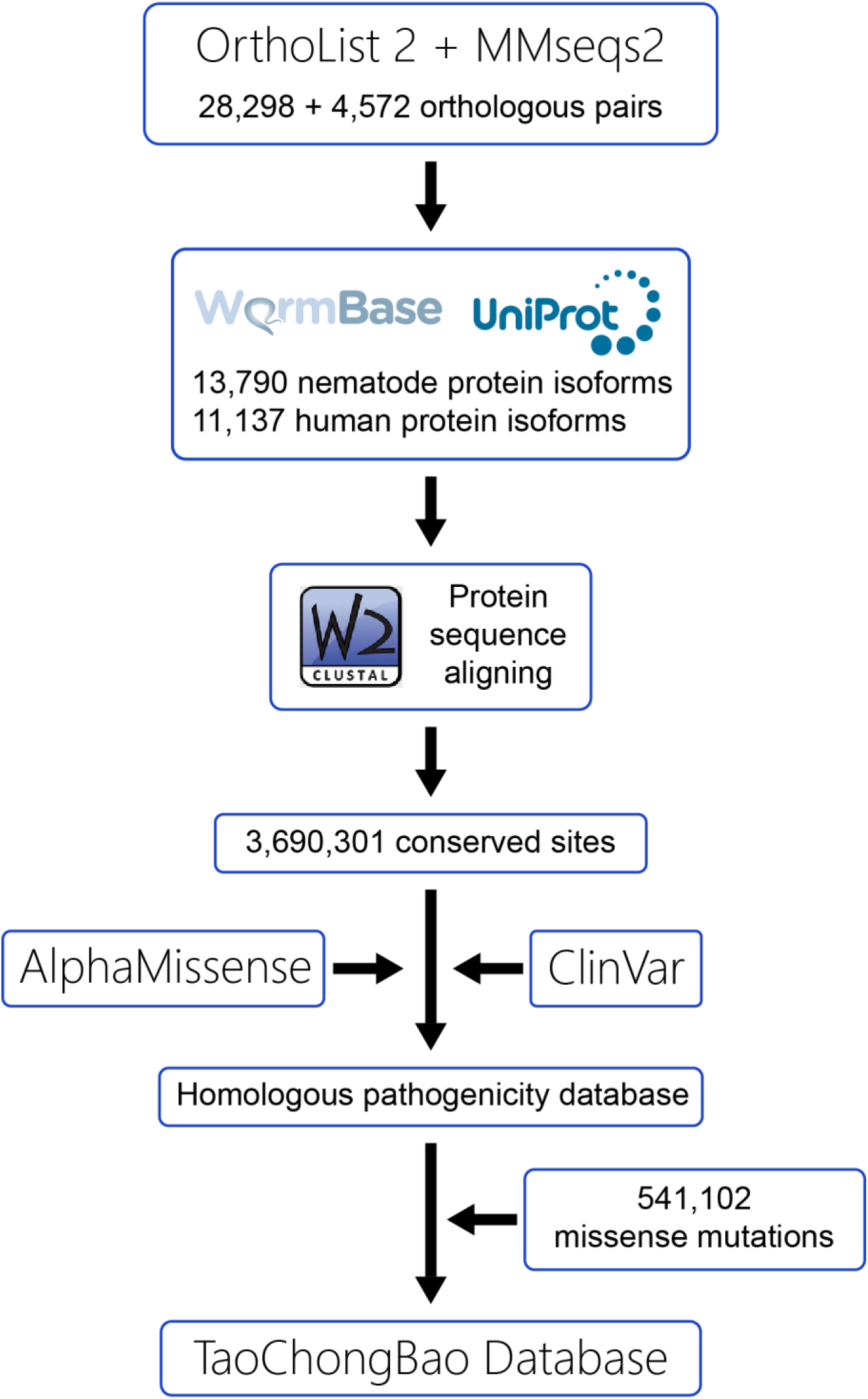
Workflow for constructing the TaoChongBao database. *C. elegans*-human orthologous pairs were retrieved from OrthoList 2 project and clustered by MMseqs2. Corresponding protein isoforms sequences were obtained from WormBase and UniProt, then aligned using Clustal W package, yielding 3,690,301 million conserved residues. Pathogenicity data from AlphaMissense and ClinVar were manually curated and integrated with the 541,102 missense mutations identified in our *C. elegans* mutant collection.

**Fig. 3.**
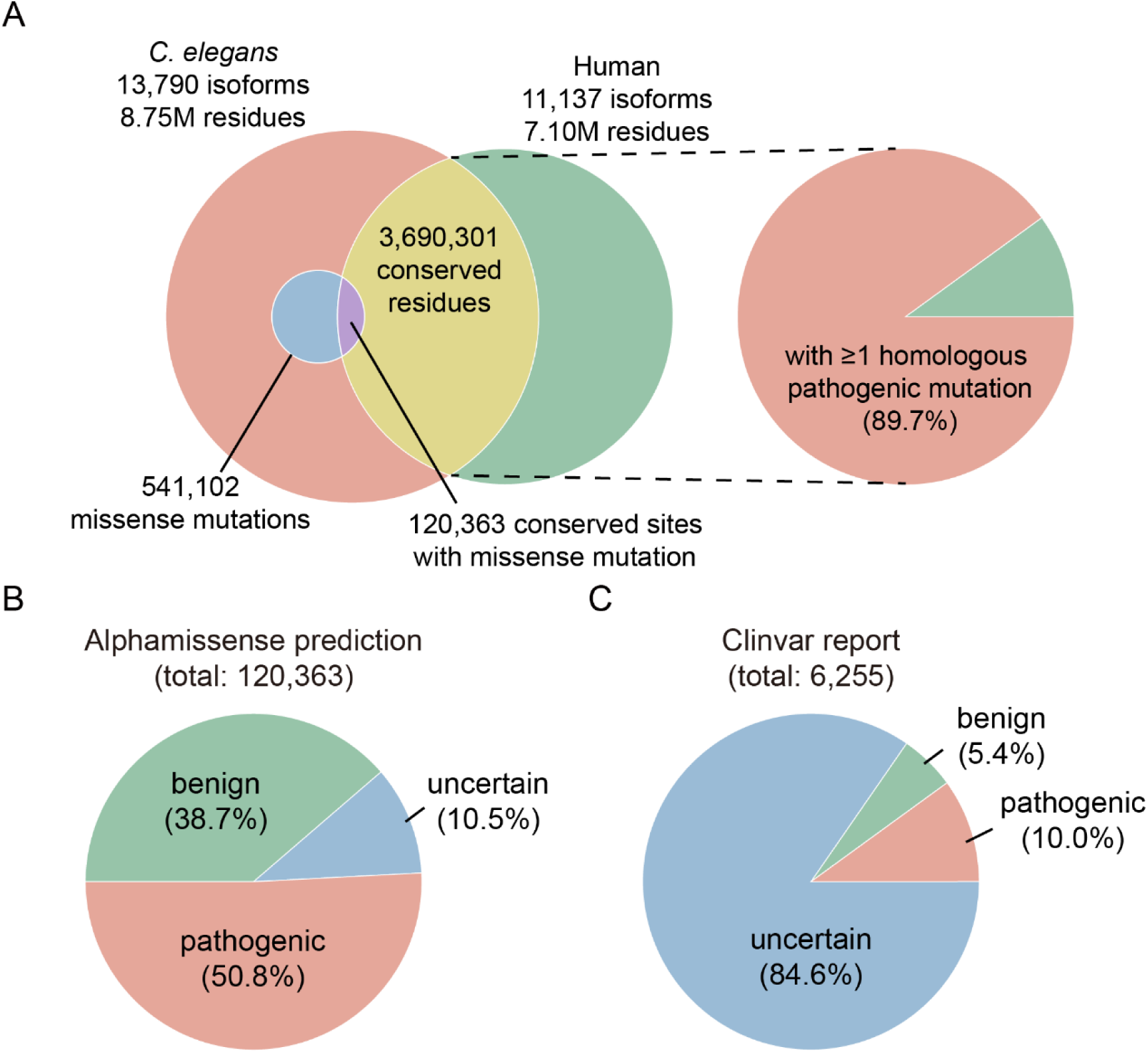
Integration of pathogenicity data and missense mutations in the *C. elegans* mutant collection. (A) Venn diagram illustrates the distribution of *C. elegans*-human conserved residues and the 541,102 missense mutations across the *C. elegans* mutant collection. (B) The predicted pathogenicity of the missense mutations occurring at conserved sites. (C) The reported pathogenicity of the missense mutations occurring at conserved sites.

Next, we integrated predicted pathogenicity data from the AlphaMissense (2), which encompasses about 71 million potential single amino acid substitutions, together with clinically reported variants from ClinVar (4). From these sources, we extracted information on substituted amino acids, predicted or reported pathogenicity, associated diseases, and other relevant annotations. We defined “homologous pathogenic mutation” as a missense mutation occurring at a conserved residue where the altered amino acid is predicted or reported to be pathogenic or likely pathogenic. Notably, our analysis revealed that 89.7% conserved sites contained at least one such homologous pathogenic mutation (Fig. 3A).

Further, we structured our sequencing results into a unified database containing essential information such as strain name, phenotypic data, gene, transcript, mutation type, position, DNA variant, and resulting protein changes. To integrate these data seamlessly with the pathogenicity annotations described above, we introduced a new field to indicate whether each mutation is classified as a homologous pathogenic mutation. From the 541,102 unique missense mutations across our *C. elegans* mutant collection, we found 120,363 (22.2%) mutations occurring at the conserved site (Fig. 3A). Approximately half of them is predicted to be pathogenic by AlphaMissense (Fig. 3B), and 5.2% (6,255 mutations) are recorded in ClinVar (Fig. 3C). Furthermore, 625 mutations are classified as pathogenic or likely pathogenic based on actual clinical observations, underscoring their potential value for the study of human genetic diseases.

Our database therefore supports two major retrieval functions: (a) users can query whether a given missense mutation in *C. elegans* corresponds to a homologous pathogenic mutation in human and, if so, obtain associated pathogenicity and disease information; (b) users can identify whether any *C. elegans* strain in our strain resource library carries homologous pathogenic mutations of interest.

### The TaoChongBao website

The TaoChongBao website interface is designed to be intuitive and user-friendly. Users can search for either a *C. elegans* or human gene of interest by entering a gene symbol, UniProt identifier, WormBase ID (for *C. elegans*), or HGNC ID (for human) (Fig. 4A). The system will automatically recognize the type of input and queries the appropriate dataset. For example, submitting a *C. elegans* gene like “*unc-104*” returns the results shown in Fig. 3B. Basic information about “*unc-104*” is displayed in the title bar, including organism, gene symbol, a brief description, and cross-references to other relevant databases (Fig. 4B ①). Since some *C. elegans* genes encode multiple isoforms, a dropdown menu lists all available isoforms, albeit in most cases only one isoform is returned (Fig. 4B ②). When multiple isoforms exist, they are sorted by the number of available strains carrying mutations in that isoform, with the first entry selected by default. User may also manually choose another isoform. This isoform-selection step is unnecessary when querying a human gene, as only one canonical isoform is included in the database.

**Fig. 4.**
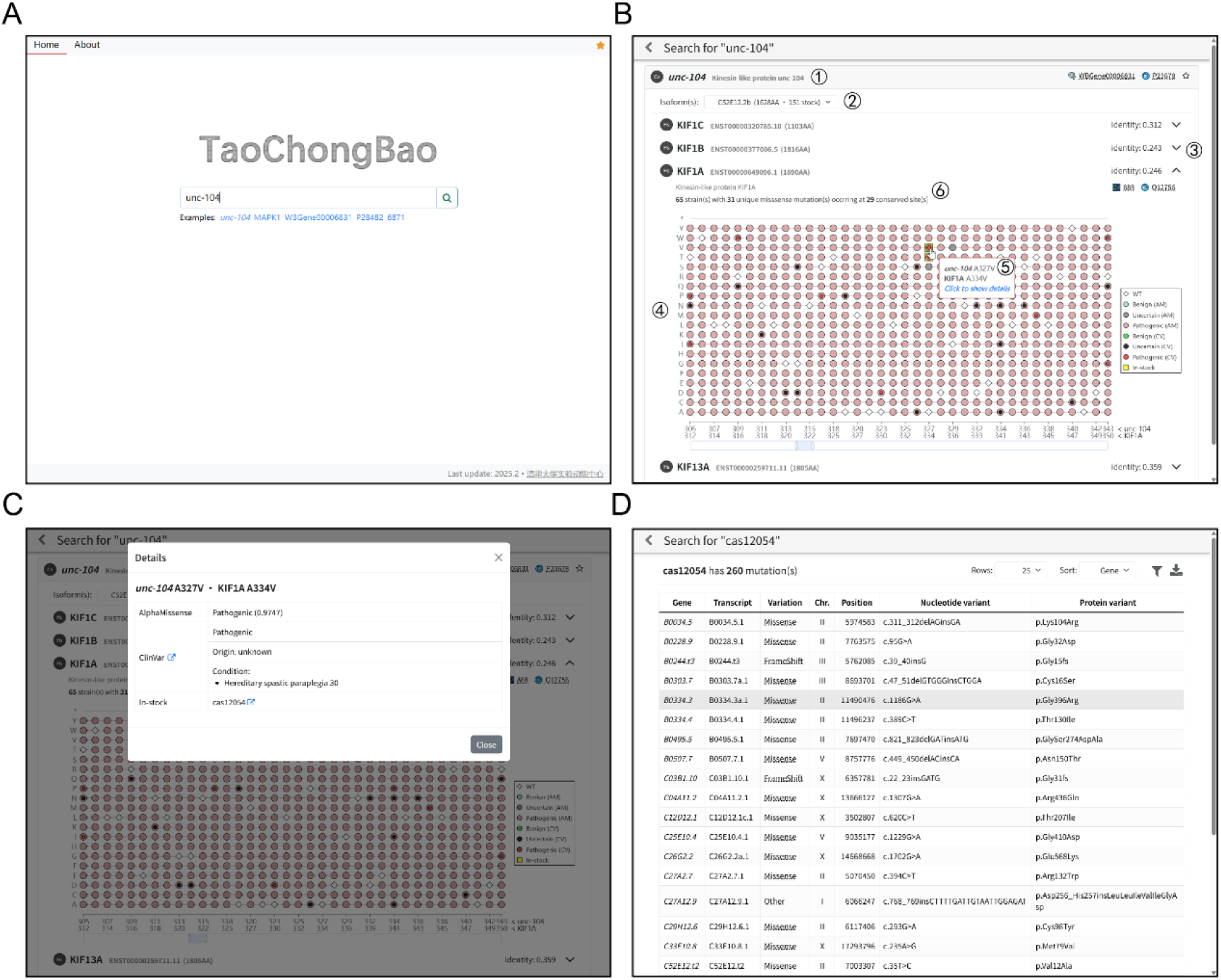
Interfaces of TaoChongBao website. (A) Home page, allowing user to search for *C. elegans* or human gene with various identifier. (B) Search results page displaying basic information for queried gene and the homologous pathogenicity information between the orthologous pairs. WT, wild-type; AM, AlphaMissense; CV, ClinVar. (C) Detail panel showing the full annotations from AlphaMissense and ClinVar, and the strain stock. (D) *C. elegans* strain page summarizing all mutations carried by the selected strain.

After selecting the desired isoform, each human ortholog is displayed as expandable tabs in the main content area (Fig. 4B ③). Each tab features a dot plot that visualizes the pathogenicity of all conserved sites between the orthologous pairs, for instance, *unc-104* and KIF1A, as shown in Fig. 4B ④. The plot includes a draggable horizontal axis indicating the conserved positions in *C. elegans* and human protein sequences, and a vertical axis indicating the amino acid at each site (* for stop codon). Wild-type residues are denoted by diamond markers, while mutated residues appear as round markers. The color of each round markers reflects the AlphaMissense predictions: red for pathogenic, green for benign, and gray for uncertain. Mutations recorded in ClinVar are indicated by a smaller dot centered on the round marker, following the similar color scheme. If a variant is present in our strain resource library, the round marker will be outlined by a yellow square. Hovering over any marker displays location and amino acid information of the corresponding variant (Fig. 4B ⑤), and clicking on it opens a panel with full annotations from AlphaMissense, ClinVar, and the strain stock (Fig. 4C). If one or more strains are available in our resource library, a jump button next to the strain name allows user to access to a page summarizing all mutations carried by that strain (Fig. 4D). Additionally, each ortholog tab also includes basic information similar to that in the title bar, along with a simple statistics summarizing the available strains harboring mutations at these conserved sites (Fig. 4B ⑥).

## Discussion

We report the largest EMS mutagenesis and whole-genome sequencing effort in *C. elegans* to date, generating a comprehensive resource of missense variants across more than 12,069 strains. This dataset includes over 20 million genomic variants, including 541,102 protein-altering missense mutations distributed across 20,914 genes—representing a twenty-fold expansion over the previous Million Mutation Project (13). To enable broad and effective use of these data, we developed the TaoChongBao database, an integrated platform which supports visualization and querying of mutations, assessment of conservation with human pathogenic residues, and access to viable strains for functional assays. By combining large-scale experimental mutagenesis with both predictive and clinical annotated pathogenicity information, this resource bridges in silico pathogenicity predictions with in vivo functional validation.

EMS mutagenesis remains one of the most cost-effective and scalable methods for generating genome-wide variant diversity. In our collection, we identified more than 120,363 missense mutations in *C. elegans* that are homologous to human mutations, including 625 that correspond to clinically annotated pathogenic alleles. For comparison, generating even a single *C. elegans* strain via genome editing approaches (e.g. CRISPR-Cas9) typically requires roughly one months and a minimum cost of US$1,000. Modeling all 625 pathogenic missense variants in our benchmark set would cost US$625,000 using targeted genome editing, and modeling the full set of 120,363 homologous human variants would exceed US$120 million—an impractical undertaking. Although EMS mutagenesis is random and cannot guarantee any specific substitution, scaling to thousands of sequenced strains substantially increases the likelihood of capturing many clinically relevant residues. Notably, our entire 12,069 strain collection was generated and sequenced for ∼US$240,000, with each strain costing about US$20 (27). These comparisons underscore that large-scale chemical mutagenesis, while complementary rather than alternative to precise editing, offers an exceptionally cost-effective strategy for generating broad allelic series that would be prohibitively expensive to obtain through genome engineering alone, thereby providing a practical platform for functional interpretation of human variation.

Chemical mutagenesis primarily generates G/C-to-A/T conversions, limiting codon coverage (9,11–13). Analyses of the Million Mutation Project suggest that EMS mutagenesis is influenced by nucleotide context and chromatin structure, meaning complete residue saturation has not yet been achieved (12,13). Some mutations, particularly lethal alleles, remain undetected, which can be addressed through temperature-sensitive or conditional genetic screens (28–30). Nonetheless, this scalable strategy provides a framework for systematically generating and functionally characterizing missense variants, with potential applications in other model organisms and human cell lines amenable to forward genetic screens.

Our strain resource library serves as a versatile tool for diverse applications. It provides genetically diverse strains that allow researchers to explore the relationships between protein sequence and function, including studies on protein folding, stability, and interactions, which are fundamental to understanding molecular biology. The library also facilitates disease modeling, enabling functional investigation of mutations analogous to human variants in a controlled setting, which can inform mechanisms of pathogenesis and aid in identifying therapeutic targets. Additionally, the resource supports suppressor and enhancer screens, allowing identification of genetic interactions that mitigate the effects of deleterious mutations.

Storing and distributing more than ten thousand WGS-sequenced strains poses a significant logistical challenge. Existing centralized repositories, such as the *C. elegans* Genetics Center or commercial platforms like Addgene, face inherent capacity limits and cannot expand indefinitely as the number of unique strains continues to grow. A more scalable solution is a decentralized, marketplace-style model—analogous to Amazon—where individual laboratories contribute their own sequenced strains, metadata, and annotations, and the broader community can browse, query, and obtain variants of interest. TaoChongBao adopts this decentralized framework, enabling laboratories to share resources directly while reducing redundancy, distributing storage burdens, and promoting a sustainable, community-driven system for genetic research.

In conclusion, TaoChongBao provides a comprehensive, community-accessible resource of *C. elegans* missense variants, integrating whole-genome sequencing data with predictive and clinical annotations. The database allows users to visualize mutations, assess conservation with human pathogenic residues, and access viable strains for functional assays. By expanding mutation coverage over twenty-fold relative to previous resources, TaoChongBao enables systematic studies of protein function, genetic interactions, and human disease modeling. Its scalable, cost-effective, and decentralized design promotes sustainable sharing of genetic resources across laboratories. We anticipate that TaoChongBao will become a widely used platform for functional residuomics, variant interpretation, and experimental genetics in both basic and translational research.

## Materials and Methods

### *C. elegans* strain culture

*C. elegans* strains were maintained as described previously (10), on nematode growth medium (NGM) plates with OP50 feeder bacteria at 20℃. All animal experiments were performed following governmental and institutional guidelines.

### EMS mutagenesis

EMS mutagenesis was performed as described before with some modifications (31,32). Worms at late L4 stage were cultured on regular NGM plates seeded with OP50 feeder till the population was just starved. Then worms from 10-20 plates were collected with M9 buffer (3 g KH_2_PO_4_, 6 g Na_2_HPO_4_, 5 g NaCl, 1 ml 1 M MgSO_4_, H_2_O to 1 L), and incubated with 50 mM EMS buffer at room temperature with continuous rotation for 4 hours. After incubation, worms were washed with M9 buffer and dispersed to 200-400 OP50-seeded NGM plates. After 20 hours, adult worms were subjected to a bleaching protocol to isolate their eggs (F1 generation). These eggs were then transferred to individual NGM plates at a density of about 10 eggs/plate and cultured at 20℃. When the F2 progeny reached the young adult stage, a single mutant was isolated under a stereomicroscope and transferred to a new NGM plate for screening, for example, ivermectin screening or dumpy/uncoordinated screening. Worm strains that stably transmitted the phenotype to the F3 generation were subjected to WGS using Illumina next-generation sequencer.

### Whole-genome sequencing

Raw WGS reads (FASTQ file) were quality-controlled using FastQC (v0.12.1), trimmed for adapters and low-quality bases by Trim_galore (v0.6.8), and aligned to WBcel235 of the *C. elegans* genome (www.ensembl.org) by BWA-MEM2 (v2.2.1). All datasets achieved >15×coverage.

After removing putative PCR duplicates using Picard (v2.27.3), variants, including single nucleotide variants (SNVs), insertion/deletion variants (indels), and multi-nucleotide polymorphisms (MNPs) were called using freebayes (v1.3.6), and annotated with SnpEff (v5.0d). To obtain homozygous SNVs, variants were filtered using vcflib tools (v1.0.3) with the criteria “QUAL > 20 & DP > 5 & AF > 0.8”, retaining locations with at least 5x coverage, ≥80% consensus variant allele, and a minimum mapping quality of 20. Essential information for constructing the TaoChongBao database was extracted using a homemade Python script.

## Data availability

All TaoChongBao resources are accessible through the web portal (https://oulab.life.tsinghua.edu.cn/homopatho/index.php).

## Acknowledgements

This work was supported by the following funding programs: the National Key R&D Program of China Grants 2024YFA1307301, 2022YFA1302700, 2019YFA0508401; the National Natural Science Foundation of China Grants 92254306, 31991190, 32270773, 32470730, 32070706, 32270721, 32430026, and 32400610; and the Postdoctoral Fellowship Program of China Postdoctoral Science Foundation GZC20240861. The genetic screens were performed at the 2022-2025 Artificial Evolution Summer School supported by the Tsinghua University and Qingshan Lake Science and Technology City in Hangzhou, China.

